# Transcriptional signatures of fentanyl use in the mouse ventral tegmental area

**DOI:** 10.1101/2023.12.18.572172

**Authors:** Megan E Fox, Annalisa Montemarano, Alexandria E Ostman, Mahashweta Basu, Brian Herb, Seth A. Ament, Logan D. Fox

**Affiliations:** Department of Anesthesiology and Perioperative Medicine, The Pennsylvania State University College of Medicine, Hershey, PA, USA; Department of Pharmacology, The Pennsylvania State University College of Medicine, Hershey, PA, USA; Institute for Genome Sciences, University of Maryland School of Medicine, Baltimore MD, USA

**Keywords:** fentanyl, self-administration, ventral tegmental area, single nuclei RNAseq

## Abstract

Synthetic opioids such as fentanyl contribute to the vast majority of opioid-related overdose deaths, but fentanyl use remains broadly understudied. Like other substances with misuse potential, opioids cause lasting molecular adaptations to brain reward circuits, including neurons in the ventral tegmental area (VTA). The VTA contains numerous cell types that play diverse roles in opioid use and relapse, however it is unknown how fentanyl experience alters the transcriptional landscape in specific subtypes. Here, we performed single nuclei RNA sequencing to study transcriptional programs in fentanyl experienced mice. Male and female C57/BL6 mice self-administered intravenous fentanyl (1.5 µg/kg/infusion) or saline for 10 days. After 24 hr abstinence, VTA nuclei were isolated and prepared for sequencing on the 10X platform. We identified different patterns of gene expression across cell types. In dopamine neurons, we found enrichment of genes involved in growth hormone signaling. In dopamine-glutamate-GABA combinatorial neurons, and some GABA neurons, we found enrichment of genes involved in Pi3k-Akt signaling. In glutamate neurons, we found enrichment of genes involved in cholinergic signaling. We identified transcriptional regulators for the differentially expressed genes in each neuron cluster, including downregulation of transcriptional repressor Bcl6, and upregulation of Wnt signaling partner Tcf4. We also compared the fentanyl-induced gene expression changes identified in mouse VTA with a published rat dataset in bulk VTA, and found overlap in genes related to GABAergic signaling and extracellular matrix interaction. Together, we provide a comprehensive picture of how fentanyl self-administration alters the transcriptional landscape of the mouse VTA, that serves for the foundation for future mechanistic studies.

## Introduction

Drug overdose deaths are at an all-time high in North America, and over the last several years, the primary cause of opioid overdose has shifted from heroin to synthetic opioids.^1^ Increasingly, illicit and counterfeit drugs contain fentanyl^2^, a potent synthetic opioid, which has led to an increase in fentanyl misuse and overdoses. Despite the increased presence of fentanyl in the drug supply, our understanding of how fentanyl alters the brain remains incomplete. Most of our mechanistic knowledge is derived from decades of work on natural and semisynthetic opioids like morphine and heroin^3^.

Opioid exposure causes lasting molecular adaptations throughout the brain, especially in mesolimbic reward regions such as the ventral tegmental area (VTA). The VTA is especially important for mediating the rewarding aspects of opioids^4,5^, and rats will self-administer intracranial morphine or fentanyl directly into the midbrain^6,7^. Acutely, opioids elevate extracellular dopamine concentrations, particularly in the downstream nucleus accumbens^8^, by two mu-opioid receptor dependent mechanisms. Activation of Gi coupled mu opioid receptors on VTA GABAergic interneurons causes hyperpolarization and subsequent disinhibition of VTA dopamine neurons. Alternatively, opioids can act directly on mu opioid receptors expressed on dopamine neurons to increase or decrease dopamine neuron firing^9^. Chronic opioid exposure also increases VTA dopamine neuron firing^10–12^, which is accompanied by transcriptional and epigenetic changes that support synaptic plasticity ^13–15^. However the VTA contains numerous neuron subtypes aside from dopaminergic, including glutamatergic, and GABAergic projection neurons, along with more recently discovered combinatorial neurons that possesses synthesis and release machinery for multiple neurotransmitters^16–18^. There is increasing evidence that both GABAergic^19^ and glutamatergic^20^ neurons in the VTA help orchestrate reward-related behaviors, and opioid exposure drives both glutamatergic and GABAergic synaptic plasticity in the VTA^4,21^. However, most studies investigating the molecular adaptations in VTA relied on whole VTA homogenates, and it is difficult to attribute changes in bulk gene expression to a specific neuronal subtype. This is further exacerbated by a lack of true neuronal subtype marker, since tyrosine hydroxylase, the rate limiting enzyme for dopamine synthesis and traditional dopamine neuron marker, is expressed by multiple neuron populations ^22–24^. VTA homogenates also contain numerous glial cell types which may transcriptionally respond to opioids and contribute to the behavioral effects as recently demonstrated in other brain regions ^25–27^. Understanding the changes that happen in specific cell-types is important, as we recently showed fentanyl exerts cell-type specific effects on neurons in the nucleus accumbens that can be targeted to alleviate opioid abstinence induced negative affect^28^.

Here, we address this gap and ask how fentanyl alters the transcriptional landscape in different ventral tegmental area cell types. We used single nuclei RNA sequencing in male and female mice that self-administered intravenous fentanyl. We integrated our data with the rat transcriptional atlas^18^ to separate into known cell types, and found divergent transcriptional responses dependent on cell type. We further compared our findings in mouse with those from a recently published bulk VTA RNAseq study using a food vs fentanyl choice procedure in rats^29^. Together, this work highlights the similarities and differences across cell type and species, and lays the foundation for subsequent mechanistic investigations.

## Methods

### Experimental Subjects

All experiments were approved by the Institutional Animal Care and Use Committee at the Pennsylvania State University College of Medicine (PSUCOM) or University of Maryland School of Medicine (UMSOM) and performed in accordance with NIH guidelines for the use of laboratory animals. Mice were given food and water *ad libitum* and housed in the PSUCOM or UMSOM vivarium on a 12:12 h light: dark cycle with lights on at 07:00. Mice were 8-9 week old male and female C57BL/6 mice bred at PSUCOM or UMSOM with original breeding pairs obtained from Jackson Labs. All mice were pair housed in corn-cob bedding, provided with nestlets, and separated by a perforated acrylic divider.

### Intravenous Surgery

At 7-8 weeks of age, mice were anesthetized with ketamine (100 mg/kg) and xylazine (12 mg/kg) and implanted with long-term indwelling jugular catheters constructed from Micro Renathane tubing (Braintree Scientific, Braintree, MA) connected to a 24 gauge cannula (P1 Technologies, Roanoke, VA) as in our previous work^30^. Mice were flushed daily with 30 µl of heparinized saline containing enrofloxacin (400 IU/mL heparin, 0.227% enrofloxacin), and allowed to recover from surgery for >5 days

### Intravenous fentanyl self-administration

One day prior to the start of self-administration, mice were habituated to the operant chambers for 30 minutes. Mice then underwent 10 consecutive days of fentanyl or saline self-administration. The first five days were under a fixed-ratio 1 (FR1) schedule of reinforcement, followed by 5 days under FR2 (3 hours/day, 1.5 µg/kg/infusion). Operant chambers (MED Associates, Saint Albans, VT) had two nose poke holes on one wall, a cue-light above each nose poke, and a house-light in the middle of the opposite wall. At the start of the session, the house-light and active nose poke were illuminated. Responses in the active nose poke triggered a 10 µL fentanyl infusion over 1 second, turned off the house-light, the active nose-poke light, and illuminated the cue light above the active nose-poke. Any additional active responses during the 1 second infusion period were recorded but did not result in further infusions. Responses on the inactive nose poke were recorded but were without programmed consequences. No prior training was used for FR1 acquisition. Mice used for single nuclei RNA sequencing were trained at UMSOM. Mice used for qRT-PCR validation were trained at PSUCOM, and also underwent a seeking test 24 hr after the last self-administration. The 1 hour seeking test was performed under extinction conditions in which responses in the active nose poke resulted in cue presentations but no drug delivery. All behavioral testing was done during the light cycle between 08:00-12:00.

### Tissue collection

Mice were euthanized by cervical dislocation and brains were rapidly removed and chilled in ice cold PBS. Cold brains were cut into 1 mm coronal sections, and one tissue punch containing VTA (14 gauge) was flash frozen on dry ice and stored at –80 until processing.

### Nuclei isolation, library preparation, and sequencing

VTA punches were thawed in ice-cold lysis buffer containing (in mM) 320 Sucrose, 5 CaCl3, 3 Mg(Ace)2, 0.1 EDTA, 10 Tris-HCl, 1 DTT, and 0.1% Triton X-100. Tissue from 4 mice per sex per drug was combined. Tissue was homogenized on ice in 500 uL lysis buffer in a 1 mL glass dounce homogenizer. Homogenates were centrifuged at 4C for 5 minutes (1700 rcf), the supernatant removed, and the pellet resuspended in 1 mL lysis buffer. Samples were passed through a 40 um cell strainer, centrifuged for an additional 5 min at 4C (500 rcf) for 5 minutes, then aspirated and resuspended in 500 uL of DPBS. A dummy sample containing equivalent tissue from naïve mice was stained with 7-aminoactinomycin D, and was used to set the gating for fluorescence activated cell sorting for additional nuclei purification (BD Aria II, 100µm nozzle, BD Biosciences). After FACS, Single-nuclei RNA-seq libraries were prepared using the 10x Genomics Chromium Next GEM Single Cell 3′ RNA v3 protocol, targeting 7,000 cells per sample. Nuclei were captured across 4 GEM wells (2 wells per sex), and each well contained nuclei from 4 separate male or female animals from either saline or fentanyl condition. Libraries were then sequenced on an Illumina Novaseq S2 flow cell (100 base pair, paired end reads) at the Institute for Genome Sciences Core at UMSOM.

### snRNA-seq analysis

A CellRanger reference package was generated using the Ensembl mm10 genome. CellRanger (v 5.0.1) filtered outputs were analyzed with Seurat v 4.0.0 in R^31^. We recovered an estimated 9093 nuclei, which were sequenced to an average read depth of 260,000 reads per nucleus (>80% saturation). We detected a median of 2800 genes per nucleus. Data were filtered to include only nuclei with >200 and <15000 genes, and <10% of reads mapping to the mitochondrial genome. We removed doublets with scDblFinder^32^, and data were then normalized using SCTransform^33^. To integrate our data from mouse VTA with the data from the rat transcriptional atlas,^18^ we applied an identical filtering, doublet removal, and normalization strategy for the raw rat data (GEO GSE168156). We then combined mouse and rat data, which were then log-normalized with a scaling factor of 10,000, and 2000 highly-variable genes was identified with FindVariableFeatures, using the vst selection method. Data were then integrated using FindIntegrationAnchors and IntegrateData using 30 principal components. Uniform approximation and projection (UMAPs) were generated following data integration using 4 resolution values (0.1, 0.2, 0.5, 1.2), and 0.5 was chosen for subsequent analysis based on clustering into known cell types. We next looked for differentially expressed genes in the mouse subset as a function of drug, using FindMarkers. To examine changes specifically in neuron subtypes, we removed non-neuronal cells, reintegrated and clustered with 30 principal components, and 3 resolution values (0.1, 0.2, 0.5). We selected resolution of 0.5 for subsequent analysis based on clustering into known neuron types. Neuron subtypes capable of co-release were determined as described in Phillips et al^18^. We examined differentially expressed genes in the mouse neuron subtypes with Libra^34^, using a pseudobulk approach that implements edgeR with a likelihood ratio test (LRT) null hypothesis framework. We generated lists of differentially expressed genes with p<0.05 in each neuron sub-cluster, and performed gene set enrichment analysis using Metascape^35^. We identified predicted transcriptional regulators of differentially expressed genes using the iRegulon^36^ Plugin in Cytoscape. Code is available at github.com/mfoxlab.

### Bulk RNA extraction and quantitative RT-PCR

RNA was extracted using Trizol (Invitrogen) and the RNeasy Mini Kit with a DNAse step (Qiagen). RNA concentration was measured on a Nanodrop (ND-8000, Thermo), and 400 ng cDNA was synthesized using reverse transcriptase iScript cDNA synthesis kit (Bio-Rad). mRNA expression changes were measured with quantitative polymerase chain reaction (qPCR) with PerfeCTa SYBR Green FastMix (QuantaBio). Quantification was performed using the –ddCt method, using glyceraldehyde 3 phosphate dehydrogenase (GAPDH) as a housekeeping gene. Primer sequences are in Table 1.

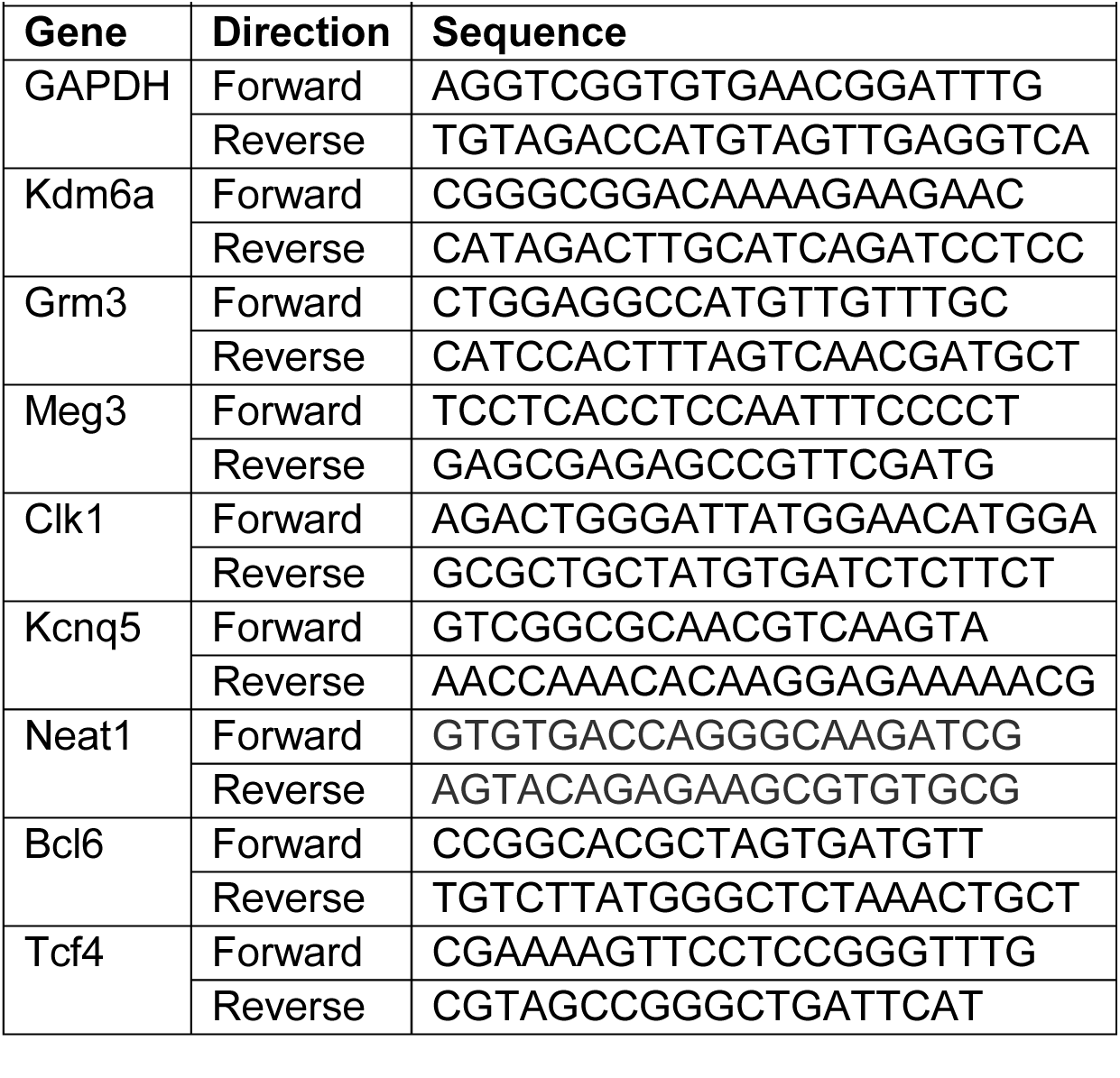
Table 1:

### Statistics

Behavioral data were analyzed using 2 or 3-way repeated measures ANOVAs in Graph Pad Prism (v 10) and JASP(jaspstats.org). Since there were no significant effects of sex, data were collapsed across sex. For seeking responses and qPCR, data were analyzed with unpaired, two-tailed t-tests.

## Results

To determine how fentanyl experience alters the transcriptional landscape in the ventral tegmental area, we trained male and female mice to self-administer intravenous fentanyl or saline on a fixed ratio (FR) 1, then a FR-2 schedule over 10 days (Timeline in **Fig 1A**). Mice self-administering fentanyl learned to discriminate between the active and inactive nose-poke. (**Fig 1B**, 3-way RM-ANOVA, day x nose-poke, F_9,108_ =3.0, p= 0.03, day x nose-poke x sex, p=0.421). Mice self-administering saline did not discriminate and had similar responses on the inactive and active nose-poke (**Fig 1C** 3-way RM-ANOVA, day x nose-poke, F_9,108_=0.684 p =0.7, day x sex x nose-poke p= 0.96). There were no significant effects of sex on the number of infusions earned (3-way RM-ANOVA, Day x Drug x Sex F_9,108_ = 0.6 p=0.78), and there were no significant differences in total fentanyl intake across the sexes (female average total intake 0.043 mg/kg, male 0.36 mg/kg, 2-way RM-ANOVA, day x sex F_9,54_ = 1.1, p=0.38). We thus combined sexes for subsequent analysis (**Fig 1D**, 2-way RM-ANOVA, Day x Drug F_9,126_ =2.8, p=0.005. Fentanyl vs Saline p<0.05 on day 7, 8, 10, Holm-Sidak).

**Figure 1.**
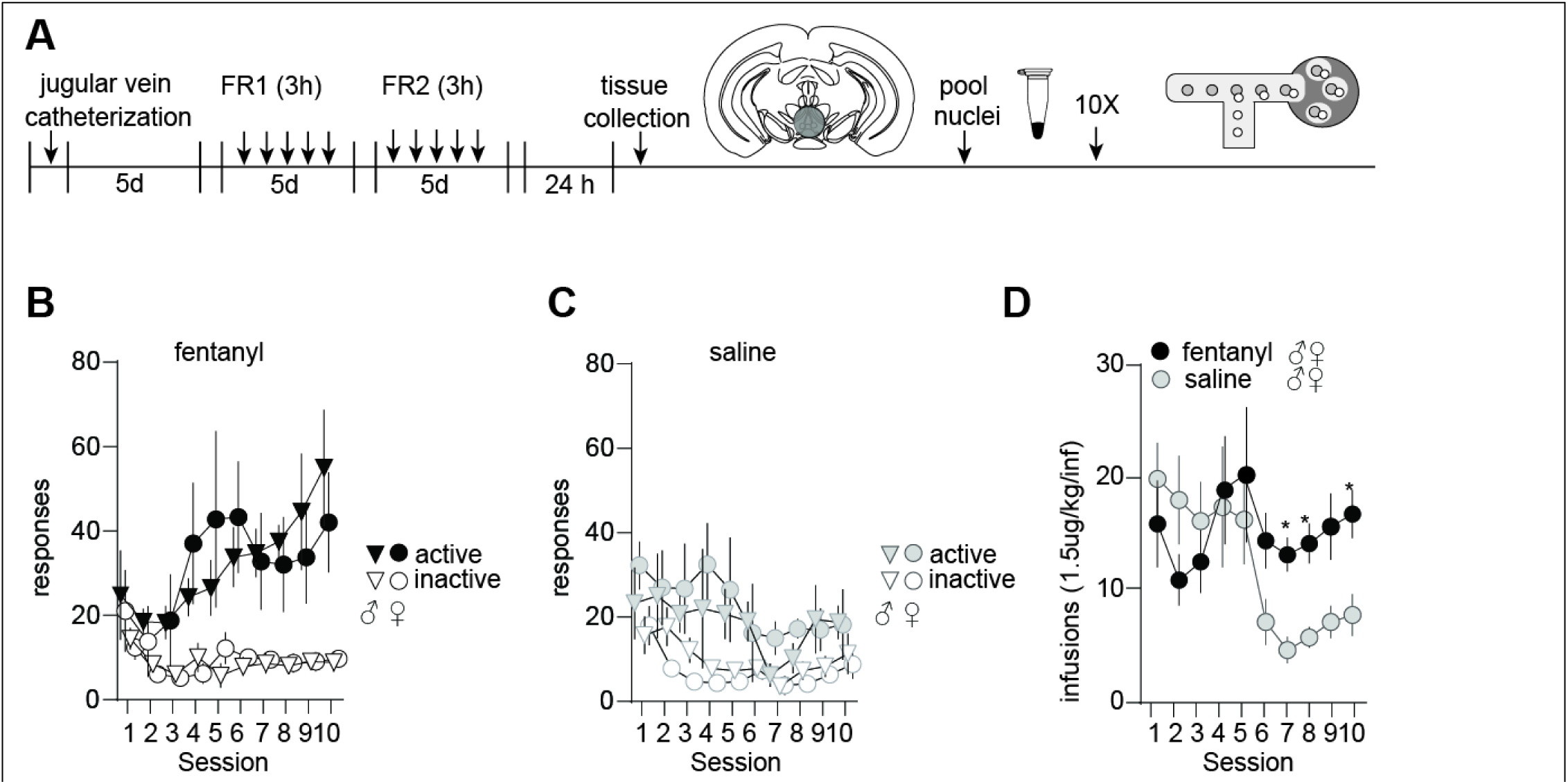
(**A**) Experimental timeline. Male and female mice (n=4/group) underwent surgery for indwelling jugular vein catheter placement. Following recovery, mice administered fentanyl (1.5μg/kg/10 μL infusion) or saline (10 μL) on a fixed-ratio (FR)-1 schedule for 5 days, then on a FR-2 schedule for 5 days. 24 hours after the last self-administration session, brains were removed and fresh tissue punches containing the ventral tegmental area (VTA) were flash frozen, then nuclei isolated and prepared for single nuclei sequencing on the 10X platform. (**B**) Responses on the active and inactive nose-poke in male (triangles) and female(circles) mice self-administering fentanyl or (**C**) saline. (**D**) Number of fentanyl or saline infusions earned collapsed across sex. *, p<0.05 fentanyl vs saline, Holm-Sidak posthoc after 2-way RM-ANOVA.

After 10 days of self-administration, we collected fresh tissue punches containing the ventral tegmental area 24 hr after the last self-administration session. We pooled dissociated nuclei based on sex and drug group, then prepared libraries for single nuclei RNA sequencing (snRNAseq) on the 10X platform. We captured a similar number of nuclei per sample across sex (**Supplemental Fig 1**). We took two approaches to look at changes in gene expression in fentanyl experienced mice. First, using the FindMarkers function in Seurat^33^, we looked for differentially expressed genes collapsed across all nuclei, in an attempt to mimic a bulk RNAseq experiment. We found few transcripts that reached a cut-off of >± 0.6 log2 fold change and adjusted p<0.05 (Red labeled genes in **Fig 2A**).

**Figure 2.**
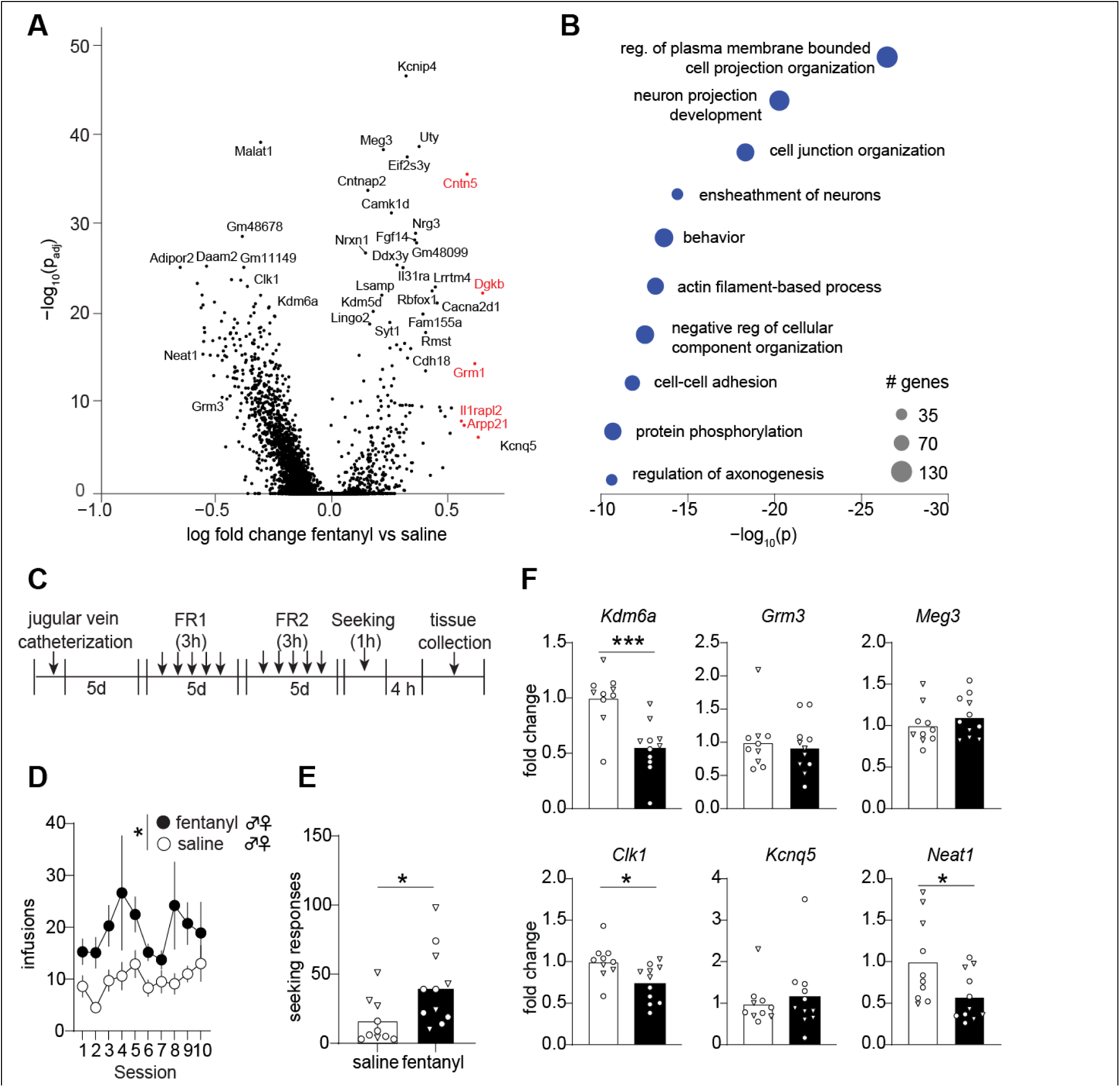
(**A**) Volcano plot showing differentially expressed genes as determined with the Seurat FindMarkers function, collapsed across cell type. (**B**) Top 10 biological process gene ontology terms identified with Metascape. (**C**) Timeline for replication cohort. After recovering from jugular catheter surgery, mice (4-6/sex/group) self-administered fentanyl or saline for 10 days. On day 11, mice were tested for drug seeking under extinction conditions in a 1 hour test. 4h after drug seeking, fresh VTA tissue punches were collected for subsequent bulk RNA extraction. (D) Number of infusions earned by male and female fentanyl or saline self-administering mice. Drug, F _1,19_= 6.6, p=0.018. (**E**) Number of responses in the active nose poke during the seeking test. t_1,19_ = 2.5, p =0.02. (**F**) Fold change expression relative to GAPDH for selected gene targets. Kdm6a t_1,19_ = 4.38 p=0.0003; Clk1 t_1,19_ = 2.71 p= 0.014; Neat1 t_1,19_=2.35, p = 0.02. Triangles denote datapoints from male mice and circles from female.

We thus used all transcripts with an adjusted p <0.05, for gene set enrichment analysis, with no log2 fold change cut-off. We performed Gene Ontology (GO) analysis with Metascape^35^, and found overrepresentation of GO terms related to neuronal structure, cell-adhesion, and protein phosphorylation (Top 10 GO terms in **Fig 2B**, all annotations in **Supplemental File 1**; gene lists in **Supplemental File 2**).

We next sought to confirm changes in bulk gene expression in an independent cohort of mice with qRT-PCR (Timeline in **Fig 2C**). In the replication cohort, mice self-administering fentanyl earned more infusions compared with saline mice (**Fig 2D**, 2-way RM ANOVA, Drug, F _1,19_= 6.6, p=0.018) and exhibited more drug seeking behavior based on active responses during a non-reinforced seeking test (**Fig 2E**, t_1,19_ = 2.5, p =0.02). We selected several genes for validation, and found that, similar to our findings with snRNASeq, fentanyl experienced mice have downregulation of the lysine demethylase *Kdm6a*, the kinase *Clk1*, and the long non coding RNA *Neat1* (**Fig 2F**). Increased expression of *Meg3* and *Kcnq5*, and decreased expression of *Grm3* failed to replicate with this approach.

To look at changes in gene expression across specific cell types, we next performed Uniform approximation and projection (UMAP) clustering after integrating our data from mouse VTA with the 21,600 rat VTA nuclei data from the transcriptional atlas^18^ (**Supplemental Fig 2**). To examine changes happening specifically in neuron subtypes, we removed non-neuronal cell types and re-clustered the rat and mouse data. (**Figure 3**). We separated the clusters into similar neurotransmitter identity clusters as in the transcriptional atlas by looking at co-expression of synthesis and transport genes, as well as previously identified markers *Gch1* and *Slc26a7* (selected clusters in **Fig 3C**, all others in **Supplemental Fig 3**. Cluster marker genes in **Supplemental File 2**). We chose four clusters for further analysis, namely cluster 16 dopamine neurons, cluster 18 dopamine-glutamate-GABA neurons (“combinatorial neurons”), cluster 13 glutamate neurons, and cluster 9 GABA neurons. We selected the latter two clusters based on low co-expression of GABAergic and glutamatergic markers, respectively. We looked at changes in gene expression in each of these sub-clusters using Libra^34^. In cluster 16 dopamine neurons, we found 407 upregulated and 403 downregulated genes in fentanyl-experienced mice relative to saline (**Figure 4A, Supplemental File 2**). In cluster 18 combinatorial neurons, we found 381 downregulated, and 361 upregulated genes (**Figure 4B; Supplemental File 2**). We identified 44 shared differentially expressed genes between dopamine cluster 16 and combinatorial cluster 18, 13 of which exhibited opposite changes in expression. We next performed gene set enrichment analysis in each cluster list with Metascape^35^. 565 differentially expressed genes in cluster 16 dopamine neurons were assigned to 8 KEGG pathways (top 3 pathways and associated genes in **Fig 4D**, all annotations in **Supplemental File 1**), which indicated fentanyl altered expression of genes involved in “growth hormone action,” “calcium signaling,” and “cAMP signaling,” among others. The top GO term for cluster 16 dopamine neurons was “transmembrane receptor protein tyrosine kinase signaling pathway.” 535 differentially expressed genes in cluster 18 combinatorial neurons were assigned to 5 KEGG pathways (top 3 in **Fig 4E**, all in **Supplemental File 1**) which indicated fentanyl altered expression of genes involved in “RNA polymerase activity” and “phosphoinositide 3-kinase/ protein kinase B activity (Pi3k-Akt)”. There was also enrichment for “hypertrophic cardiomyopathy’, which contains genes important for cell adhesion. The top GO term for cluster 18 combinatorial neurons was “DNA damage response.” 708 genes in Cluster 9 GABA neurons were assigned to 4 KEGG pathway terms (**Supplemental File 1**). Like dopamine neurons, this included “PI3K-AKT signaling.” There was also enrichment for “Neuroactive ligand receptor interaction”,” axon guidance,” and the “Hippo signaling pathway,” which modulates cell survival. The top GO term for cluster 9 GABA neurons was “regulation of nervous system development.” 884 differentially expressed genes in cluster 13 glutamate neurons were assigned to 9 KEGG pathway terms, among them the notable “morphine addiction”, “cholinergic synapse”, and “adrenergic signaling.” The top GO term for cluster 13 glutamate was “monoatomic ion transmembrane transport” (**Supplemental File 1**).

**Figure 3.**
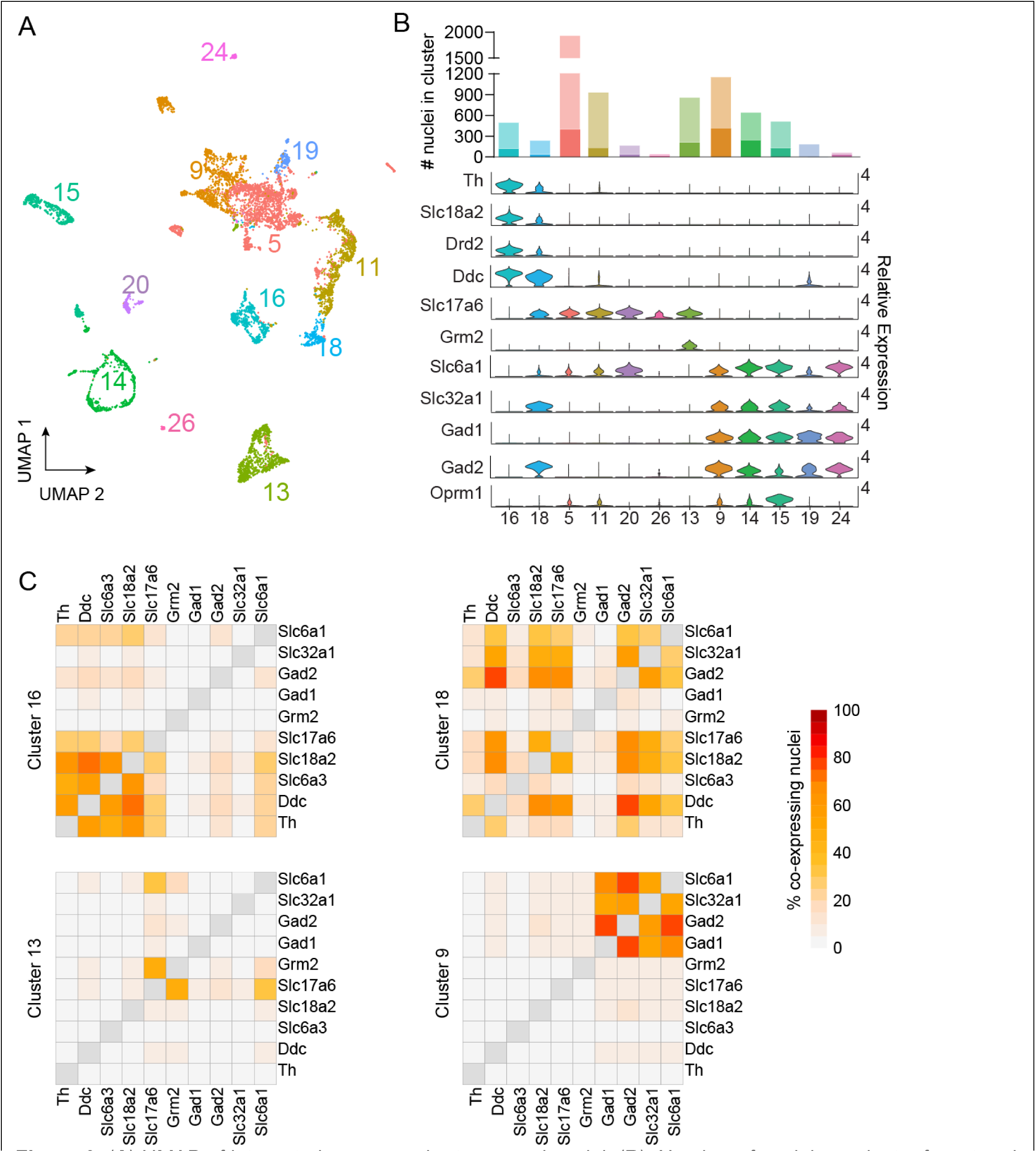
(**A**) UMAP of integrated mouse and rat neuronal nuclei. (**B**). Number of nuclei per cluster for rat and mouse data. Violin plots showing enrichment of cell type markers. Dopaminergic markers: Th, tyrosine hydroxylase, Slc18a2, vesicular monoamine transporter, Drd2, dopamine D2 receptor, Ddc, dopa decarboxylase, Glutamatergic markers, Slc17a6, vesicular glutamate transporter, Grm2, metabotropic glutamate receptor 2,GABAergic markers, Slc6a1, GABA transporter protein type 1 (GAT1), Slc32a1, vesicular GABA transporter, Gad1, glutamate decarboxylase 1, Gad2, glutamate decarboxylase 2. Oprm1, mu opioid receptor. (**C**) Heatmaps of nuclei co-expressing genes involved in dopamine, glutamate, and GABA synthesis and transport.

**Figure 4.**
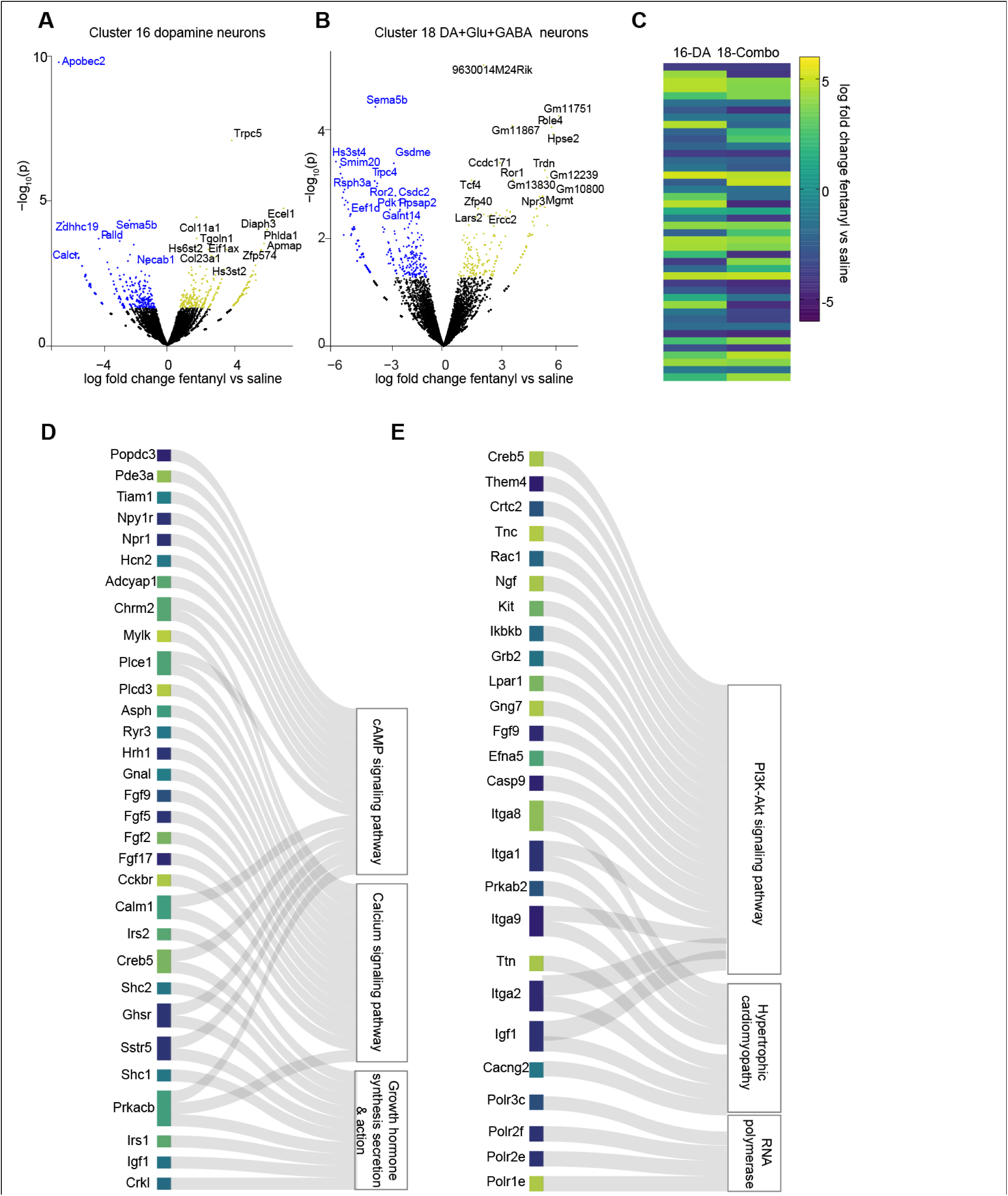
(**A**) Volcano plot showing differentially expressed genes in Cluster 16 dopamine neurons and (**B**) Cluster 18 dopamine-glutamate-GABA combinatorial neurons. Downregulated genes are in blue and upregulated genes are in yellow (**C**) Heatmap showing log fold change expression for the 44 shared differentially expressed genes between cluster 16 dopamine and cluster 18 combinatorial neurons. (D) Sankey plot showing the top 3 KEGG pathway terms for cluster 16 and (**E**) cluster 18 differentially expressed genes, the genes that are contained in that enriched pathway, and the log fold change expression in fentanyl vs saline within that cluster is encoded in color.

We next took the list of differentially expressed genes in each cluster and looked for common transcription factor binding sites upstream of the gene promoters using iRegulon^36^ (Top 5 regulators for the selected clusters in **Figure 5A-D**, all in **Supplemental File 1**). We compared the lists of transcriptional regulators with the lists of differentially expressed genes in our selected neuron clusters, and selected differentially expressed transcriptional regulators for qRT-PCR replication. Although not statistically significant, we found a trend towards similar decreased expression of cluster 9 regulator *Bcl6*, and increased expression of cluster 18 regulator *Tcf4* (*Bcl6*: t_1,13_=1.7, p=0.1, *Tcf4*, t_1,15_=2.0, p=0.06, **Figure 5E**). We next looked for concordance in the patterns of expression of the transcription factor target genes by counting the number of differentially expressed target genes within the corresponding cluster (Example network of *Bcl6* target genes in cluster 9 **in Figure 5F**).

**Figure 5.**
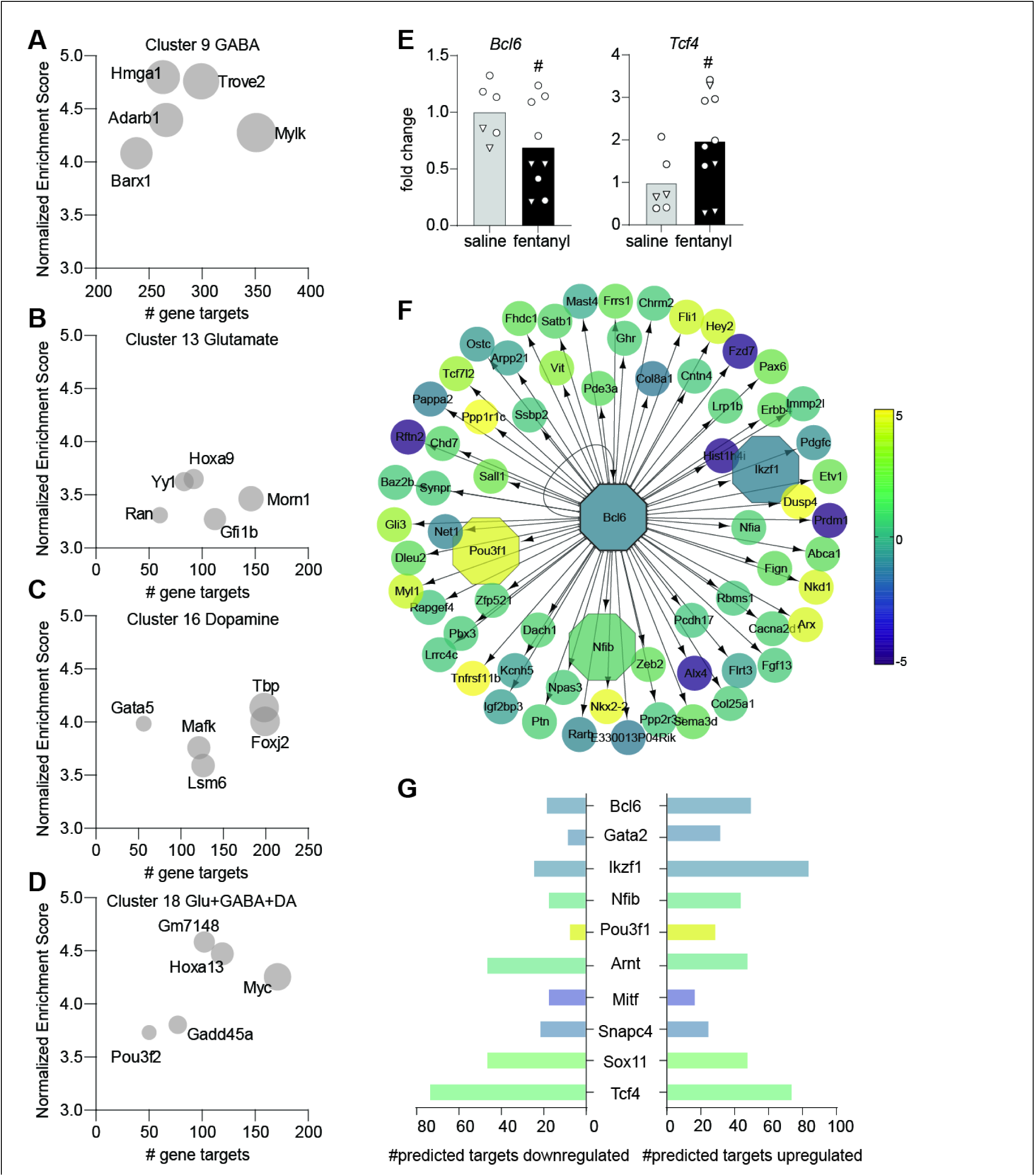
Top five predicted transcriptional regulators of differentially expressed genes in (**A**) Cluster 9 GABA, (**B**) Cluster 13 Glutamate, (**C**) Cluster 16 dopamine, and (**D**) Cluster 18 combinatorial neurons. The iRegulon Normalized Enrichment score is on the Y-axis, and the number of predicted gene targets on the X axis. (**E**) Fold change expression relative to GAPDH for selected predicted transcription factors in bulk VTA. Bcl6: t_1,13_=1.7, p=0.1, Tcf4, t1,15=2.0, p=0.06. (**F**) Network of genes regulated by Cluster 9 regulator Bcl6. Log fold change fentanyl vs saline within Cluster 9 is encoded in color. (**G**) All predicted transcriptional regulators also differentially expressed in snSeq, and the number of their targets that are downregulated or upregulated within that cluster. Bcl6, Gata2, Ikzf1, Nfib, and Pou3f1 are Cluster 9 regulators, the remainder are Cluster 18. The log fold change fentanyl vs saline of that transcriptional regulator is encoded in color, using the same scale as (F).

Most patterns were concordant with the expected direction, i.e. increased expression of *Pou3f1*, a positive regulator of transcription, was linked with a greater number of gene targets upregulated in cluster 9; decreased expression of transcriptional repressor *Bcl6*, was linked with a greater number of gene targets upregulated in cluster 9. Many of the predicted transcriptional regulators, including *Tcf4*, can both promote and suppress transcription. Increased expression of *Arnt*, *Sox11*, *Tcf4*, or decreased expression of *Mitf*, which have context dependent transcriptional regulatory effects, were associated with a mostly equivalent number up and downregulated target genes in cluster 18 (**Figure 5G**).

We next compared the genes identified in our mouse snSeq dataset with a recently published bulk RNAseq dataset from fentanyl self-administering rats^29^. We compared our differentially expressed genes to both the male and female rat VTA gene lists using two approaches. We first used the list we generated in Figure 2A, using the FindMarkers function and adjusted p<0.05, and compared these genes to those reported as p<0.05 differentially expressed in male or female rats (**Figure 6A**, **left**). This resulted in only 11 species-shared upregulated, and 49 downregulated genes, with the remainder of overlapping genes exhibiting opposite changes. We next compared our cluster specific lists determined by pseudo-bulking (**Figure 6A, right**). This resulted in 124 species-shared upregulated, and 83 downregulated. For the common upregulated and common downregulated genes, we performed gene set enrichment analysis with Metascape^35^. The common upregulated genes were assigned to five KEGG pathways (top 3 shown in **Fig 6B**), including “GABAergic synapse,” “Neuroactive ligand receptor interaction,” and “ferroptosis.” The common downregulated genes were assigned to 7 KEGG pathways (top 3 in **Fig 6C**), including extracellular matrix –receptor interaction, focal adhesion, and dilated cardiomyopathy, the latter pathway containing genes important for calcium signaling. The common up and downregulated genes between fentanyl experienced mouse and rat were spread across mouse cell type clusters. The top GO terms for rat and mouse common genes were all related to development (upregulated: “positive regulation of neuroblast proliferation, positive regulation of neurogenesis”, downregulated: “head development”, “brain development.”)

**Figure 6.**
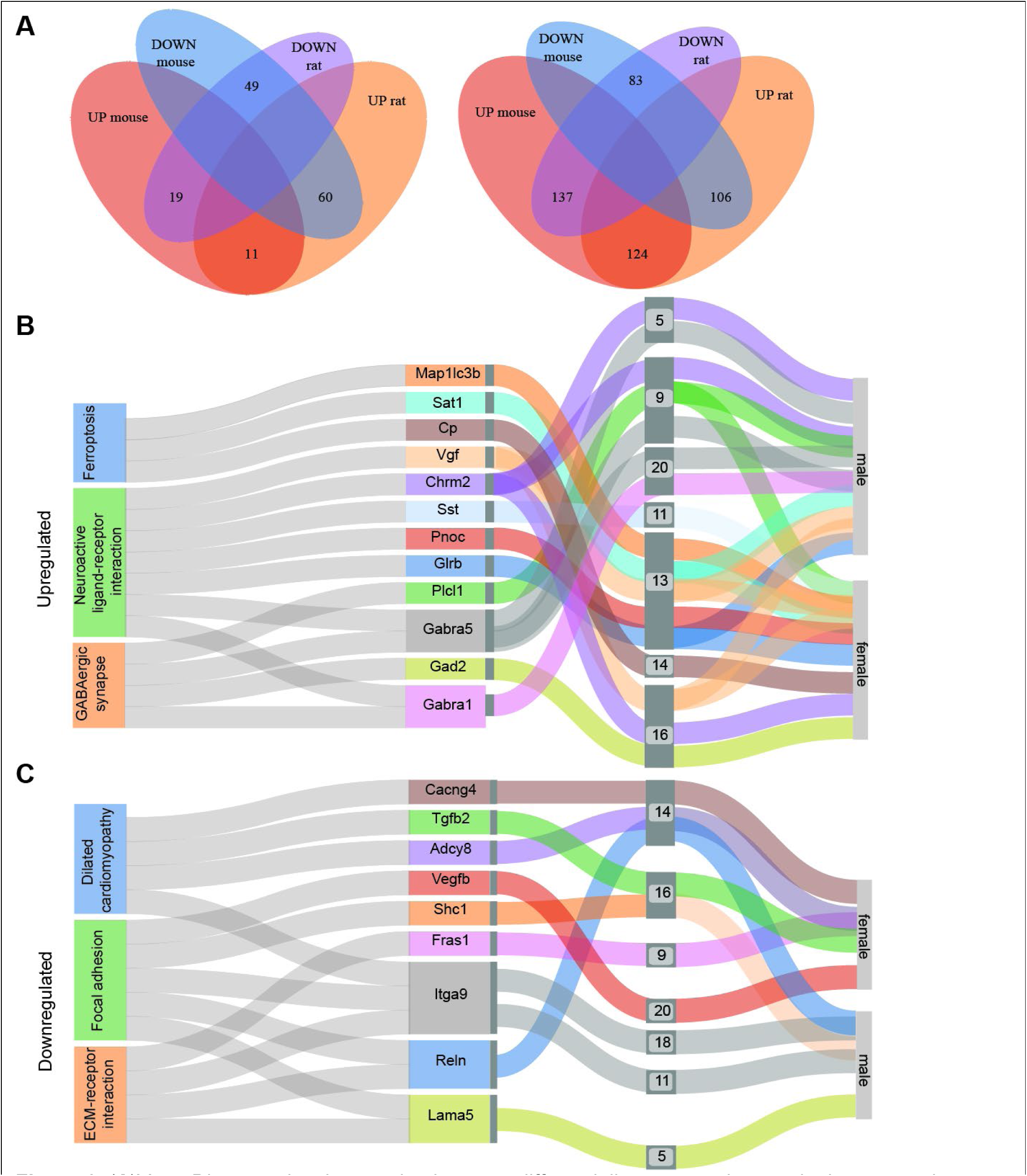
(**A**)Venn Diagram showing overlap between differentially expressed genes in the mouse dataset in this paper, and the rat data in Townsend et al(ref) with mouse gene lists taken from the FindMarkers padj<0.05 on the left and lists from pseudobulking with Libra p<0.05 on the right. Sankey plots showing Top 3 KEGG pathway terms in the mouse cluster-specific and rat common upregulated genes (B) and (**C**) downregulated genes. The common genes are shown as belonging to numbered mouse neuron clusters, and if they were differentially expressed in male or female rats.

## Discussion

Here, we identified different transcriptional responses to fentanyl self-administration in the mouse VTA. Collapsed across cell-type, we found downregulation of *Kdm6a*, *Clk1*, and *Neat1* that replicated with bulk qRT-PCR. We used a transcriptional atlas to separate cells into previously described types, and found many cell-type specific transcriptional programs that were altered by fentanyl exposure. The cell type specific changes were, in turn regulated by different transcription factors. Finally, we showed many genes are differentially expressed in both fentanyl experienced rats and mice using a recently published dataset. Together, this work provides a detailed look into the transcriptional landscape of the fentanyl experienced ventral tegmental area, and will serve as a launching point for many future mechanistic investigations.

Our primary goal in this work was to uncover cell type specific changes in gene expression in the VTA as a function of fentanyl experience. We chose to conduct this work in mice as they are the preferential model for transgenic lines, and any identified gene targets could be directly tested in genetically defined cell types. Our mice learned to self-administer intravenous fentanyl, as evidenced by discrimination of the active vs inactive nose-poke, and a greater number of infusions earned compared with saline control. They also exhibited drug seeking behavior, as evidenced by an increased responding under extinction conditions. One limitation in our study is that the transcriptional effects here are a combination of the pharmacologic actions of fentanyl, in addition to the act of self-administration. Future studies could employ a yoked control to examine which changes in gene expression can be attributed to volitional intake vs mu opioid receptor activation.

In our initial analysis, we employed very strict cut-offs, and were left with very few genes that met criteria. One source of this is the low number of replicates in this study. We elected to pool mice from the same sex and drug condition in favor of a greater depth of sequencing as opposed to lower coverage of more samples. Despite lowering the stringency, we were still able to replicate many of the differentially expressed genes in bulk qRT-PCR. Our future work will use more cell-type specific approaches such as fluorescence in situ hybridization and translating ribosome affinity purification in Cre/Flp driver lines. Even with this additional layer of validation, the gene candidates will still have to be tested for causal roles in fentanyl relapse behavior, so we have chosen to disseminate this data in its present form so as to facilitate more expedient mechanistic inquiries.

With our more lenient cut-offs, we found several interesting patterns of gene expression changes. In dopamine neurons, we found mostly upregulation of genes associated with calcium signaling, mostly downregulation of genes associated with growth hormones, and both up– and down-regulation of genes involved in tyrosine kinase receptor signaling. Of the differentially expressed genes in these categories, decreased expression of *Igf1* (insulin-like growth factor 1) is a standout, as recent work in the prefrontal cortex showed decreased IGF1 protein in mice that self-administered oral fentanyl, and IGF-1 replacement in the PFC attenuated fentanyl seeking behavior^37^. Decreased expression of *Calcr* (calcitonin receptor) is another intriguing target, as *Calcr* expression has bidirectional effects on opioid self-administration in nucleus accumbens neuron subtypes^37^. The top upregulated genes in dopamine neurons was the extracellular matrix protein *Ecel1*, and the actin polymerization regulator *Diaph3*, both of which are implicated in synaptic function and structural remodeling. These will be interesting gene targets to explore, as chronic morphine is known to induce structural plasticity of VTA dopamine neurons^12,38,39^, and extracellular matrix proteins are an area of interest in the opioid use disorder field^40^. We found increased expression of Insulin receptor substrate *Irs1* and *Irs2*, opposite of the decreased Irs proteins after chronic morphine in rat VTA^41^. Other similarities and differences with our data in fentanyl self-administration dopamine neurons and that of chronic morphine mouse total VTA ^42^ are a concordant increases in the zinc finger protein *Bcll1b,* the kinase *Dclk3*, the postsynaptic protein *Synpo*, and a discordant decrease in the kinase *Sgk1*, synaptic vesicle protein *Synpr*, and increase in kinase *Hipk2*. Of our selected neuron clusters, cluster 9 GABA neurons had the most overlap with genes in Heller et al^42^, with concordant downregulation of transcription factor *Alx4*, and upregulation of ankyrin domain protein *Ankrd33b,* transcriptional regulators *Arx, Ddn*, *Lhx6*,, adapter protein *Baiap2*, long noncoding RNA *Dlx6os1*, deaminase *Gda*, GTPase regulators *Ngef*, *Rab40b*, and cytoskeletal regulator *Wipf3*. There was discordant decrease in kinase anchoring protein *Akap5*, cAMP phosphoprotein *Arpp21*, and discordant decreases and increase in transcription factors *Rarb*, and *Tcf712*, respectively. There were only 3 common genes in cluster 13 glutamate (concordant down actin protein *Actg2*, concordant up *Wipf3*, discordant down phosphodiesterase *Pde10a*), and 2 in cluster 18 combinatorial (discordant down *Baiap2*, concordant up G protein subunit *Gng7*). It will be important to determine if these common up-and down-regulated genes play a causal role in volitional opioid intake and relapse, as recent work showed that deleting *Sgk1* in dopamine neurons, upregulated after chronic morphine and cocaine, does not disrupt drug reward behaviors, and only a catalytically inactive mutant decreases cocaine conditioned place preference, but not self-administration ^43^.

We found some overlap in the differentially expressed genes within the two dopamine neuron populations, including concordant up-regulation potassium channel *Hcn1*, and cAMP binding protein *Creb5.* These targets are of particular interest as chronic opioids are known to disrupt expression of potassium channels ^12^, and cAMP-mediated upregulation of HCNs in dopamine neurons was recently shown to promote cocaine self-administration^44^. We also found enrichment for Pi3k-Akt signaling pathway genes in the combinatorial neurons and our selected GABA neurons cluster. This included upregulation of neurotrophic factors *Ngf* and *Gdnf*. However this upregulation in *Gdnf* is contrary to the decreased *Gdnf* mRNA in rat VTA after an equivalent abstinence period from heroin self-administration^45^. One of the most interesting findings was an upregulation of mRNA for the mu opioid receptor, *Oprm1*, in cluster 13 glutamate neurons. Recent work indicates that mu opioid receptors are in fact, expressed on VTA glutamate neurons, and that they modulate excitatory transmission onto neurons that release dopamine into the nucleus accumbens core^46^. We also found decreased expression of adrenergic receptor *Adra1b*. Activation of α1 receptors increases glutamate release onto VTA dopamine neurons^47^. Together, increased Gi signaling from mu opioid receptors and decreased signaling through α-1 receptors would reduce glutamatergic input to dopamine neurons, and may be a potential source of decreased dopamine release seen after opioid experience^48^. The numerous gene targets that control cellular excitability will be interesting candidates to pursue as the field of non-dopaminergic mechanisms in the VTA continues to expand.

To further explore the patterns of differential expression in the cell types, we identified transcription factors that have binding sites upstream of differentially expressed gene promoters, as altered transcription factor expression likely contributes to the differential patterns of gene expression within each cell type cluster. Many of the transcription factors we identified were also differentially expressed in our sequencing data, and in a direction concordant with their known regulatory effects. One of these differentially expressed transcription factors, *Sox11*, is also upregulated in the nucleus accumbens after chronic morphine^11^. Identifying the common transcriptional regulators is a useful tool in a candidate gene discovery pipeline, as it is impossible to test how every individual differentially expressed gene mediates behavior. Targeting a transcription factor allows one to manipulate expression of many different genes at once, which is a strategy we have successfully used to change transcriptional programs in the fentanyl abstinent nucleus accumbens.^28^

We also sought to identify gene expression changes that persist across multiple fentanyl self-administration models. We found a number of common differentially expressed genes in the VTA from fentanyl self-administering mice and rats^29^. Many of the commonly upregulated genes were involved in GABAergic signaling, and included two genes for GABA(A) receptor alpha subunits, *Gabra1*, *Gabra5*, and GABA synthesis enzyme *Gad2*. VTA GABA signaling is augmented by both mu opioid receptor activation and morphine withdrawal^49^, and chronic morphine produces a similar upregulation of α1 and α5 subunits in hippocampus^50^, but their role in mediating opioid relapse hasn’t yet been investigated. The top KEGG pathway for shared downregulated genes in rats and mice was extracellular matrix receptor interaction. Downregulation of the extracellular glycoprotein *Reln* is especially interesting, as global reduction in Reelin protein is associated with increased locomotor sensitization to cocaine^50^. We can find no mention of most of the common differentially expressed genes between rats and mice regarding a role in opioid related behavior, but since they are preserved across species and fentanyl intake paradigms, they warrant future follow up.

Overall, we have provided a comprehensive look at the transcriptional consequences of fentanyl use in the mouse ventral tegmental area, including diverse changes across cell types. It will be important to follow up on the top gene targets and their common transcriptional regulators in a cell type specific manner to establish a causal role for the gene targets in driving opioid use and relapse behavior. By highlighting the similarities and differences across cell type and species, we have laid the foundation for numerous future mechanistic investigations.

---

**Supplemental Figure 1.**
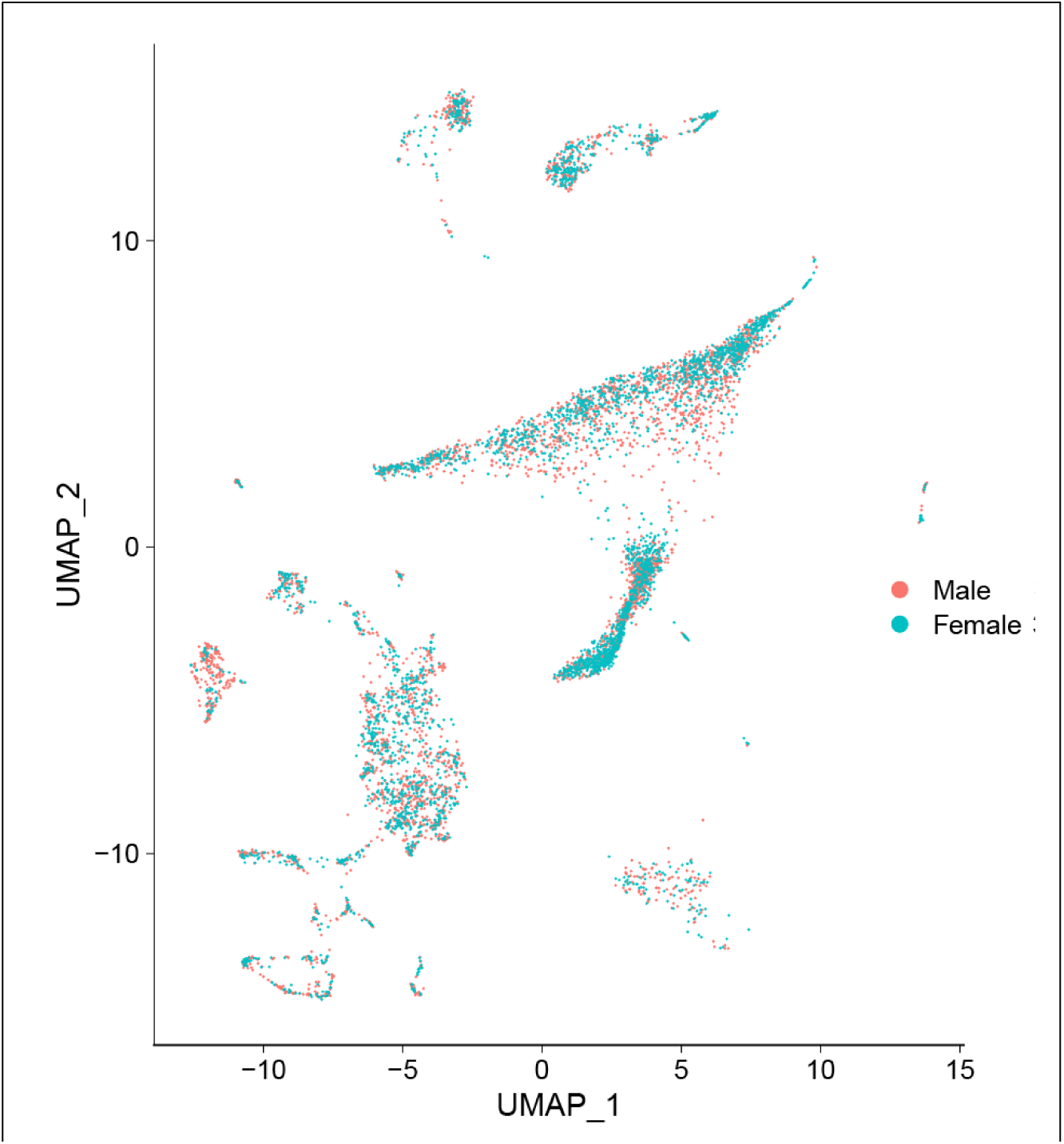
Uniform approximation and projection (UMAP) of integrated mouse nuclei, colored by sex.

**Supplemental Figure 2.**
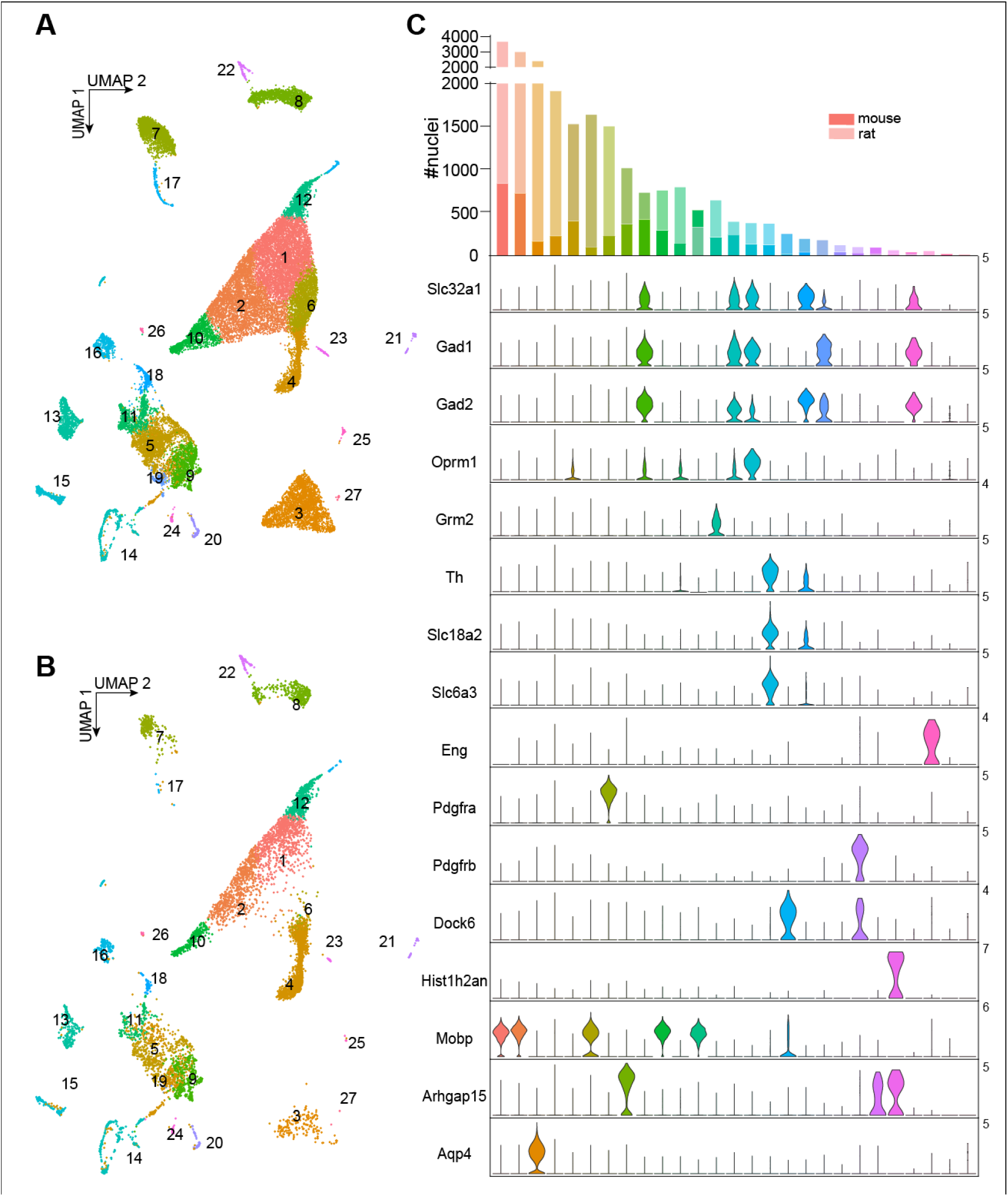
(A) Uniform approximation and projection (UMAP) of integrated mouse and rat nuclei. (B) UMAP of mouse only subset. (C) Number of nuclei per cluster for rat and mouse data. Violin plots showing enrichment of cell type markers. Neuronal markers: Slc32a1, vesicular GABA transporter, Gad, glutamate decarboxylase 1 and 2, Oprm1, mu opioid receptor, Grm2, metabotropic glutamate receptor 2. Th, tyrosine hydroxylase, Slc18a2, vesicular monoamine transporter 2, Slc6a3, dopamine transporter. Endothelial marker, Eng, endoglin. Polydendrocyte marker Pdgfra, platelet derived growth factor receptor a. Mural cell marker, Pdgfrb. Oligodendrotyce precursor markers, Dock6, dedicator of cytokinesis 6, Hist1h2an, histone cluster 1, H2an, Oligodendrocyte marker, Mobp, myelin associated oligodendrocyte basic protein; Microglia marker, Arhgap15, Rho GTPase activating protein 15, Astrocyte marker, Aqp4, aquaporin 4

**Supplemental Figure 3.**
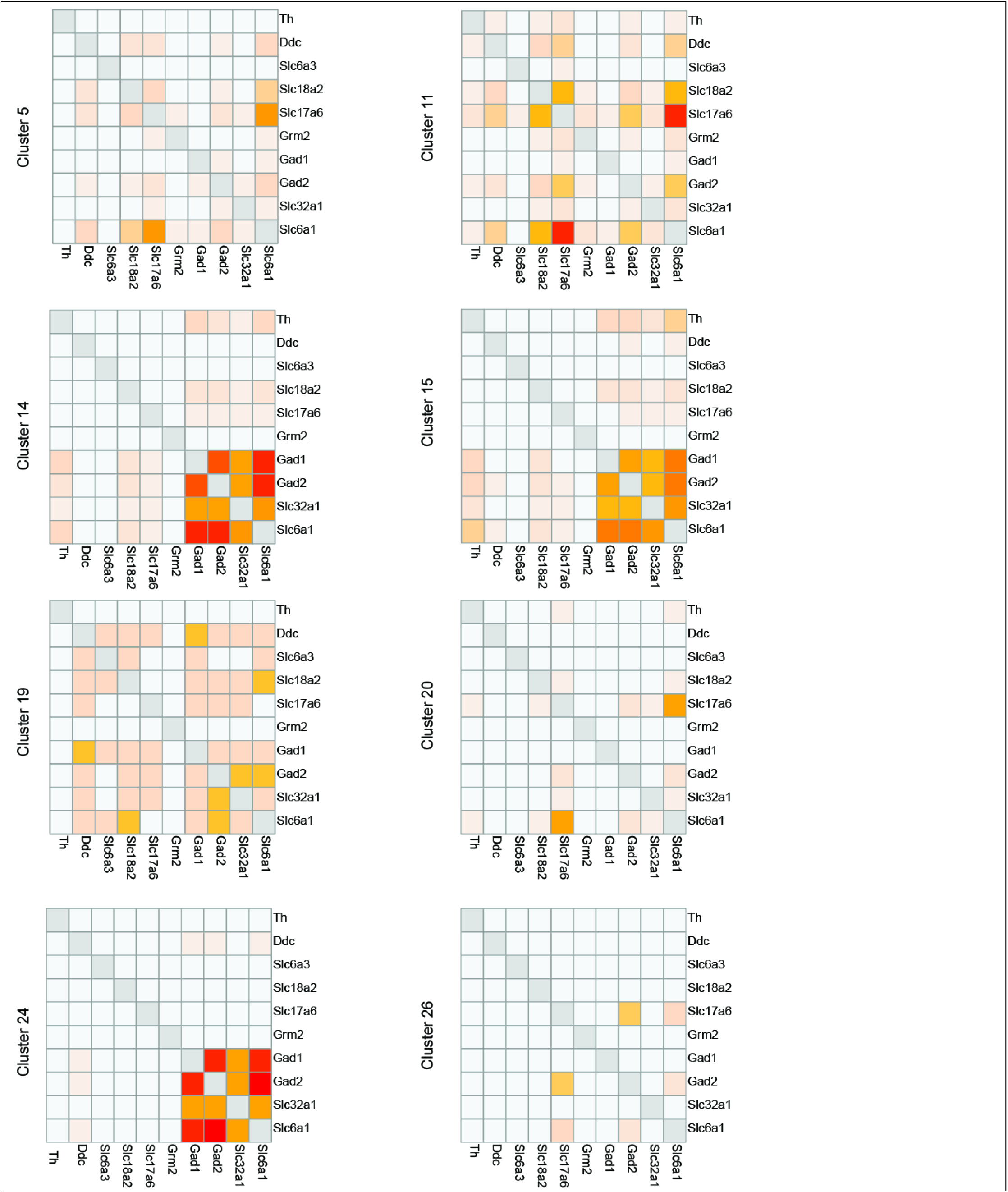
Heatmaps of nuclei clusters co-expressing genes involved in synthesis and transport of GABA, glutamate, and dopamine.

